# Ischemic lesions to inferior frontal cortex alter the dynamics of conscious visual perception

**DOI:** 10.1101/2024.08.24.609496

**Authors:** M. Fritsch, J. Michely, L. Jaeckel, I. Rangus, C. Riegler, J. Scheitz, CH Nolte, P. Sterzer, V. Weilnhammer

## Abstract

Whether the prefrontal cortex is part of the neural correlates of conscious visual perception has been subject to longstanding debate. Recent work, using functional magnetic resonance imaging (fMRI) and repetitive transcranial magnetic stimulation (rTMS) to induce virtual lesions, has illustrated a key role of the right inferior frontal cortex (IFC) in the detection and resolution of perceptual ambiguities. Here, we sought to validate an active role of the IFC in conscious perception by evaluating how loss-of-function in patients with ischemic stroke in this region influences processing of ambiguous visual information. To this end, twenty-three patients (11 female, mean age 70.65 ± 1.2 years) with chronic (>6 months), right-hemispheric ischemic stroke lesions within the MCA-territory (9 patients with IFC lesions, 14 without) performed a bistable perception task, having to report the perceived direction of rotation of an ambiguous random-dot-kinematogram (RDK). As hypothesized, patients with IFC lesions showed significantly fewer perceptual changes compared to patients without IFC lesions (IFC 44.9 ± 46.3 s, non-IFC 28.2 ± 37.1 s, T(182.9) =3.1, p = 2.2 x 10^-3^). Importantly, this effect remained significant when controlling for age, sex, stroke severity and lesion volume (T(5.97) = −2.9, p =0.026). Our results support the notion that the IFC is crucial for resolving perceptual ambiguities, suggesting an active role of frontal cortex in shaping conscious visual experience.

## Introduction

In order to adequately navigate our surroundings, we need to transform the inherently ambiguous data registered by our sensory organs, into unambiguous conscious experience (Brascamp et al., 2018). One key approach to investigate the underlying neural mechanisms is through ambiguous visual stimuli that hold equal sensory information for two mutually exclusive percepts, resulting in bistable perception (Leopold and Logothetis, 1999; Sterzer et al., 2009). It has been proposed that spontaneous changes in conscious experience during the viewing of such stimuli are caused by the unconscious accumulation of perceptual conflict during the alternating phases of perceptual stability (Hohwy et al., 2008; Weilnhammer et al., 2017). Recent work suggested a key role of the prefrontal cortex in the detection and transformation of such ambiguous conflicts into unambiguous conscious experience. Specifically, functional magnetic resonance imaging (fMRI) revealed a neural correlate of accumulating perceptual conflict in the right inferior frontal cortex (IFC) (Weilnhammer et al., 2021, 2017). Further, inducing a transient virtual lesion, by inhibiting right IFC activity with repetitive transcranial magnetic stimulation (rTMS), led to a reduction in spontaneous changes during bi-stable perception (Weilnhammer et al., 2021), thus suggesting a causal role of IFC in shaping conscious visual experience. Here, we sought to validate the role of the IFC in conscious perception by evaluating how loss-of-function due to ischemic stroke in this region would impact the resolution of perceptual conflict during bistable perception. We hypothesized that, analogous to rTMS-induced virtual lesions in IFC, patients with ischemic IFC lesions would experience fewer perceptual changes while viewing ambiguous stimuli.

## Results

### Clinical Data

Twenty-three patients with chronic ischemic lesions (mean time since stroke onset 16.2 months) in the right middle cerebral artery (MCA) territory participated in a behavioral experiment using a previously established bistable-perception task (Table 1) (Weilnhammer et al., 2021). Nine patients had MCA lesions affecting the right IFC (“IFC”) and 14 patients had lesions sparing the right IFC (“non-IFC”) (Figure 1.2). Groups did not differ in sex (female, IFC 44 ± 0.5%, non-IFC 46 ± 0.5%, p = 0.9). There were significant between-group differences in age (IFC 69 ± 9.6, non-IFC 71 ± 11.3 years, p = 0.03) and stroke severity, as measured with the National Institutes of Health Stroke Scale (NIHSS) (Brott et al., 1989), at discharge (IFC 2.0 ± 2.6, non-IFC 1.0 ± 1.3 points, p = 2.5 x 10^-7^). Further, we found a significant difference in lesion size (IFC 45154 ± 51694 voxels, non-IFC 903 ± 1137 voxels, T(143.11) = 10.282, p = 2.2 x 10^-16^).

**Table 1.**

|  | IFC | Non-IFC | p |
| --- | --- | --- | --- |
| **Clinical Data** |  |  |  |
| N | 9 | 14 |  |
| Age (y) | 69 ± 9.6 | 71 ± 11.3 | p = 0.03 |
| Sex (f, %) | 44 ± 0.5 | 46 ± 0.5 | p = 0.9 |
| NIHSS at discharge | 2.0 ± 2.6 | 1.0 ± 1.3 | p = 2.5 x 10<sup>-7</sup> |
| Lesion volume (voxel) | 45154 ± 51694.13 | 903.3 ± 1137.27 | p = 2.2 x 10<sup>-16</sup> |
| **Measures of Interest** |  |  |  |
| Phase Time (s) | 44.9 ± 46.3 | 28.2 ± 37.1 | p = 2.2 x 10<sup>-3</sup> |
| **Control Parameters** |  |  |  |
| Response Times (s) | 0.69 ± 0.18 | 0.70 ± 0.16 | p = 0.51 |
| Accuracy (%) | 88.33 ± 13.05 | 85.0 ± 17.46 | p = 0.30 |
| Uncertainty (%) | 2.8 ± 16.3 | 0.95 ± 8.2 | p = 0.14 |

**Figure 1.**
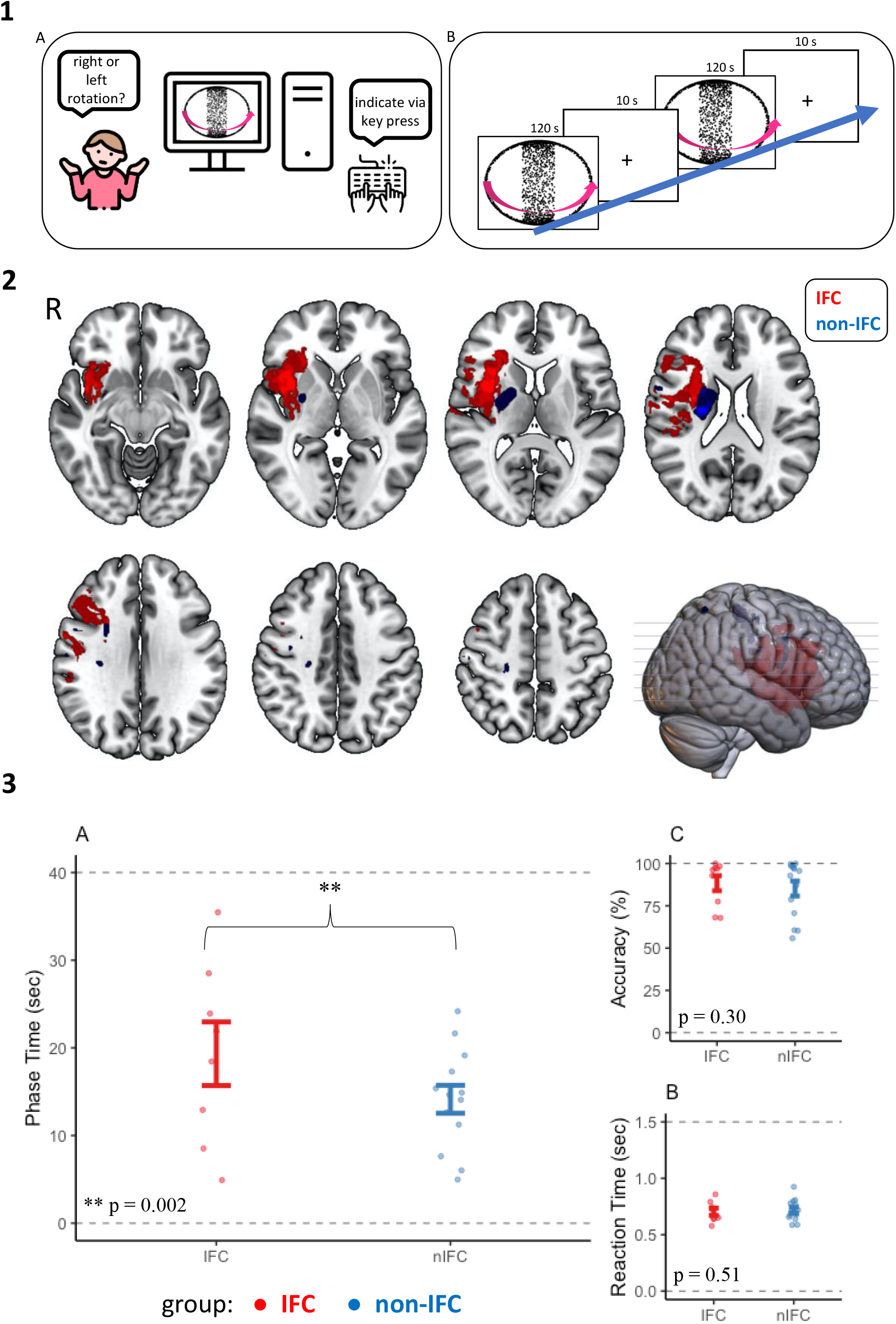

### Fewer Perceptual Changes in IFC vs. non-IFC lesions

In the behavioral experiment, participants viewed ambiguous and unambiguous versions of a random-dot-kinematogram (RDK) and indicated changes in perceived rotation direction via key presses by choosing between “right-rotation”, “left-rotation” and “unclear” (figure 1.1). The unambiguous version served as a control condition to measure task performance. In line with our hypothesis, patients with IFC lesions showed significantly longer phase durations between perceptual changes during viewing of the ambiguous RDK resulting in a lower frequency of consciously perceived changes compared to patients without IFC lesions (IFC 44.9 ± 46.3 s, non-IFC 28.2 ± 37.1 s, T(182.9) = 3.1, p = 2.2 x 10^-3^). This effect remained significant when controlling for age, sex, clinical stroke severity and lesion volume (T(5.97) = −2.9, p =0.026) (figure 1.3). Critically, average phase duration did not correlate with lesion volume across participants (rho = 0.122, p = 0.61).

### Control Parameters

We found no group differences in other performance measures, such as response times – measured in relation to an overlap within the stimulus (IFC 0.69 ± 0.18 s, non-IFC 0.70 ± 0.16 s, T(209.44) = −0.65, p = 0.51), uncertainty – the amount of perceptual states deemed “uncertain” by participants (IFC 2.8 ± 16.3 %, non-IFC 0.95 ± 8.2 %, T(183.36) = 1.49, p = 0.14), or accuracy – the amount of correctly noted perceptual changes in the unambiguous control run (IFC 88.33 ± 13.05 %, non-IFC 85.0 ± 17.46%, T(87.86) = 1.04, p = 0.30) ruling out general between-group differences in basic aspects of motivation, attention or information processing. (figure 1.3, Table 1). For a more detailed explanation of computing please see the methods section.

## Discussion

Here we show that patients with ischemic stroke lesions in the right IFC exhibit significantly longer phase durations, and therefore fewer perceptual changes, than control patients with adjacent lesions not affecting the IFC, when viewing a bistable stimulus. This finding substantiates the notion that IFC plays a causal role in the resolution of perceptual ambiguities, as previously suggested (Brascamp et al., 2018; Leopold and Logothetis, 1999; Sterzer et al., 2009). While some previous functional neuroimaging work investigating the role of frontal brain regions in bistable perception suggested that IFC merely reflects a feedforward process in response to perceptual processes occurring within sensory cortices (Brascamp et al., 2015; Frässle et al., 2014), other studies have suggested an active role of IFC in initiating spontaneous perceptual changes through feedback to sensory cortices (Sterzer and Kleinschmidt, 2007; Weilnhammer et al., 2021, 2017, 2013).

Our current finding critically expands the latter view and is in line with previous work showing that ‘virtual’ IFC lesions due to inhibitory rTMS reduce the rate of perceptual changes during bistable perception (Weilnhammer et al., 2021). The observation that ischemic IFC lesions equally reduce the rate of perceptual changes further stresses the causal importance of IFC-activity in conscious visual perception. It should be noted, though, that these observations do not refute the idea that IFC is implicated in feedforward processing. According to predictive processing accounts of bistable perception, a change from one percept to another while viewing a bistable stimulus might relate to an accumulation of prediction-errors that are elicited from visual cortex in response to the contradicting evidence of an alternative percept within the stimulus (Hohwy, 2012; Weilnhammer et al., 2017). IFC may thus have a role in both detecting and then resolving this perceptual conflict arising from visual cortex by inducing a change in perception (Weilnhammer et al., 2021). Our new finding strongly supports the notion that the signaling of perceptual conflicts in IFC is critically involved in the resolution of perceptual ambiguities.

It is important to acknowledge limitations. Firstly, our sample size is small, limiting analytical approaches. A larger sample size would allow for more sophisticated imaging methods such as voxel-based lesion symptom mapping or connectome analyses. That said, our sample is comparable in size to similar studies evaluating cognitive processes in stroke patients (Aron et al., 2003; Choo et al., 2022). Including only patients with lesions in one vascular territory, and excluding lesions other than ischemic stroke, is a strength of our sample. Our approach allows for a better comparability of groups and rules out possible secondary lesion effects such as edema in hemorrhagic lesions or tumors (Lim-Hing and Rincon, 2017). However, we must acknowledge a certain heterogeneity regarding lesion size with significant differences in lesion volume between groups. This is likely due to the fact, that stroke lesions involving more peripheral areas of the MCA-territory, such as IFC, are typically either territorial lesions (involving all of MCA-territory) or scattered, and therefore tend to be larger (Koennecke et al., 2001). This was also true for our IFC-sample. Importantly, lesion volume did not affect or correlate with our parameter of interest “phase time”, thus rendering a significant role of this group difference on our parameter of interest unlikely.

Further, there were significant between-group differences regarding age and NIHSS at discharge. Differences in age and NIHSS were small, but it could be argued that the IFC group might have shown poorer overall performance due to larger stroke burden. However, none of these parameters – NIHSS, age and lesion volume -affected our main finding when used as covariates in the statistical analyses. Further, we calculated and compared relevant parameters of task performance between groups. Neither accuracy, measured in the unambiguous control run only, nor response times differed significantly between the IFC and the non-IFC group. This suggests that the effect of a longer phase time in the IFC-group in fact relates to an inherent function of IFC and not to an overall impairment in task performance.

Overall, we were able to show that, similar to transient, rTMS-induced lesions, ischemic lesions to IFC reduce perceptual changes during the viewing of bistable stimuli. This finding emphasizes the important role of IFC in the resolution of perceptual ambiguities and thus, in determining the contents of conscious perception. Understanding how our brains generate and update the contents of consciousness has significant implications for unraveling the mechanisms behind pathophysiology, such as psychotic symptoms, that may occur when the access of conflicting sensory data to consciousness is dysregulated. Therefor addressing this question could offer profound insights into the neural foundations of consciousness itself.

## Methods

### Participants

Patients were recruited through the Stroke Unit of the Department of Neurology at Charité Campus Benjamin Franklin, Berlin, between 12/2019 and 12/2022. For comparison of patients with and without right IFC-lesions, we included only patients with acute right-hemispheric ischemic stroke within the right middle cerebral artery (MCA) territory. This ensured inclusion of IFC-lesioned patients in part of the sample and allowed a high comparability between groups. Patients with acute lesions in other vascular territories as well as with chronic lesions in the posterior territory were excluded. Further exclusion criteria were severe or uncorrected visual impairment as well as pre-existing neurological or psychiatric comorbidities. Participants gave written, informed consent. Overall, 46 patients were recruited. Out of the initially recruited patients, 16 patients could not be reached for a follow-up, or were not interested in further participation in our behavioral study. Further, 7 suffered recurrent stroke or another illness preventing further participation. Finally, 23 patients (11 female, mean age 70.65 ± 1.2 years) with chronic right-hemispheric lesions within the MCA-territory participated in our behavioral study and were included in our analyses. Stroke severity at admission and discharge was assessed using the National Institutes of Health Stroke Scale (NIHSS, (Brott et al., 1989)). Behavioral data was collected in the chronic lesion phase (≥ 6 months post-stroke, (Vaidya et al., 2019)) at the Department of Psychiatry at Charité Campus Mitte, Berlin (Table 1).

### Imaging

Participants received routine clinical stroke imaging, including diffusion-weighted imaging (DWI) and fluid-attenuated inversion recovery (FLAIR) MRI sequences. Acute lesions were diagnosed in DWI sequences with 2.5 mm slice thickness. Patients later received additional imaging at the time of behavioral assessment, consisting of FLAIR and T1 sequences, to control for recurrent stroke. Both imaging sessions were performed on 3-Tesla Siemens Magnetom Prisma scanners respectively (Siemens Medical Solutions, Erlangen, Germany). Allocation to “IFC” and “non-IFC” lesion groups was carried out by a neurologist blinded for behavioral and clinical data and not involved in data analyses using the Automated Anatomic Labeling (AAL) atlas (Tzourio-Mazoyer et al., 2002). Lesions within the right IFC were defined as being located in the area comprised of the anterior insula and the inferior frontal gyrus (pars triangularis and pars opercularis). Nine patients showed MCA-lesions affecting the right IFC (“IFC”), and 14 patients showed MCA-lesions without affecting the right IFC (“non-IFC”) (an overlap of the lesions for the IFC and non-IFC group respectively is shown in figure 1.2).

### Behavioral Experiment

Patients performed a bistable, structure-from-motion task established in prior work of our group (Weilnhammer et al., 2021). Participants were asked to report changes in the perceived rotation direction of a spherical, discontinuous RDK (figure 1.1). Perception of the spherical object was induced by the formation of random dots into two intersecting rings rotating around a vertical axis (diameter: 15.86°, rotational speed: 12 s per rotation, rotations per block: 10, individual dot size: 0.12°) (figure 1.1). Stimuli were presented on an LCD-Monitor (60 Hz, 128031024 pixels, 60 cm viewing distance, 37.82 pixels per degree visual angle) using Psychophysics Toolbox 3 (RRID:SCR_002881) and MATLAB R2019b (The MathWorks, RRID:SCR_001622). In total, participants performed 4 runs, each consisting of six blocks (120 s each) that were separated by 10 s of fixation intervals.

In the first run, a completely unambiguous version of the stimulus was presented to assess participants’ task performance. The stimulus was disambiguated by attaching a stereo disparity signal (1.8° visual angle) to all dots on the stimulus surface using dichoptic presentation with red-and-blue filter glasses (left eye: red channel, right eye: blue channel). Here, direction of rotation changed several times throughout the block by inverting the stereo disparity signal, thus creating stimulus-driven, exogenous changes in conscious experience. In all following runs, the sphere was perceptually ambiguous, with no stereo disparity cues, holding equal amount of stimulus evidence for left- and rightward rotation, thus inducing bistable perception. Participants were uninformed regarding the ambiguity of stimuli and were instructed to report changes in rotation direction via button-press on a keyboard, differentiating the rotation of the stimulus’ front-surface between i) left, or ii) right, or iii) unclear direction.

### Imaging Analyses

Lesions were manually drawn on the DWI sequences using MRIcro (https://www.nitrc.org/projects/mricro/). DWIs and the corresponding, individual lesion maps were then re-oriented, normalized and resliced using SPM 12 and the Clinical Toolbox (SPM12; Wellcome Trust Centre for Neuroimaging, University College London, London, UK), running on MATLAB (The MathWorks, Inc., Natick, MA). A T1 template originating from elderly subjects and provided by the Clinical Toolbox was used for normalization (Rorden et al., 2012). Further, lesion maps were registered to MNI standard space. Normalized lesion maps contained 2 × 2 × 2 mmisovoxels. Lesion overlay maps were created in MRIcro GL (https://www.nitrc.org/projects/mricrogl) (figure 1.2).

### Behavioral Analyses

As in prior work (Weilnhammer et al., 2021, 2017), bistable perception, marked by endogenously driven changes between the perceived direction of rotation in ambiguous runs (runs 2-4), was measured as phase-time, i.e. the elapsed time between perceptual changes, in seconds. Here, a change in conscious visual experience of the sphere only occurs when the two intersecting rings overlap (figure 1.1). Thus, timing of the perceptual events was corrected to the last overlapping configuration preceding a button-press (Weilnhammer et al., 2021). Additionally, the following basic performance measures were derived from participants’ behavioral reports: response time (measured as the time between a button press and the last preceding overlap of the intersecting rings), accuracy (measured as accurate reporting of stimulus induced perceptual changes in rotation direction during the first run), and uncertainty (measured as the amount of perceptual states marked as uncertain).

### Statistical Analyses

Statistical analyses were carried out using MATLAB and R. Two-way repeated measures analysis of variance (rm-ANOVA) and two-sided, paired t-tests were used for group comparisons. Spearman correlations were used to test correlations. We applied the R-method glm with a binomial link-function for logistic regression, using R-packages lme4 and afex for linear mixed effects modeling (RRID:SCR_0156564).

## Figure Legend

1. **Experimental Design:** (A) -subjects underwent a computer based behavioral task, reporting perceived rotation of a spherical RDK via key press. B -Subjects performed 4 runs of which the first showed a disambiguated version of the stimulus. Three runs were completely ambiguous thus inducing bistable perception. Each run consisted of 6 blocks at 120 seconds each that were separated by 10 seconds of fixation intervals. Perceived rotation was reported via button press on a keyboard.
2. **Lesion locations** of both groups respectively shown as overlay maps. All patients had lesions in the right MCA territory. Nine participants had lesions involving the right IFC – shown in red. IFC was defined as being located in the area comprised of the anterior insula and the inferior frontal gyrus (pars triangularis and pars opercularis) using the AAL atlas. 14 patients had lesions without involvement of the right IFC and served as the control group -shown in blue. R = right hemisphere.
3. **Results** of the behavioral task: (A) patients with IFC lesions showed significantly longer phase durations between perceptual changes (IFC 44.9 ± 46.3 s, non-IFC 28.2 ± 37.1 s, T(182.9) = 3.1, p = 2.2 x 10^-3^). (B + C) The control parameters “Reaction Time” and “Accuracy” – calculated in the disambiguated run – did not differ significantly between groups (response times – IFC 0.69 ± 0.18 s, non-IFC 0.70 ± 0.16 s, T(209.44) = −0.65, p = 0.51, accuracy (IFC 88.33 ± 13.05 %, non-IFC 85.0 ± 17.46%, T(87.86) = 1.04, p = 0.30).

